# Nearly unbiased estimator of *N*_e_/*N* based on kinship relationships

**DOI:** 10.1101/872168

**Authors:** Tetsuya Akita

**Affiliations:** National Research Institute of Fisheries Science, Japan Fisheries Research and Education Agency, 2-12-4 Fukuura, Kanazawa, Yokohama, Kanagawa, 236-8648, Japan

## Abstract

This study develops a nearly unbiased estimator of the ratio of the contemporary effective mother size to the census size in a population, as a proxy of the ratio of contemporary effective size to census size (*N*_e_*/N*). The proposed estimator is based on both known mother–offspring (MO) and maternal-sibling (MS) relationships observed within the same cohort, in which sampled individuals in the cohort probably share MO relationships with sampled mothers. The rationale is that the frequency of MO and MS pairs contains information regarding the contemporary effective mother size and the (mature) census size, respectively. Therefore, the estimator can be obtained only from genetic data. Moreover, We also evaluate the performance of the estimator by running an individual-based model. The results of this study provide the following: i) parameter range for satisfying the unbiasedness, and ii) guidance for sample sizes to ensure the required accuracy and precision, especially when the order of the ratio is available. Furthermore, the results demonstrate the usefulness of a sibship assignment method for genetic monitoring, providing insights for interpreting environmental and/or anthropological factors fluctuating *N*_e_*/N*, especially in the context of conservation biology and wildlife management.

## 1 INTRODUCTION

The Estimation of the ratio of the contemporary effective population size to the census size (*N*_e_*/N*) has attracted much research attention for providing information about a current population, especially in the context of conservation biology and wildlife management (Frankham, Bradshaw, & Brook, 2014; Palstra & Fraser, 2012). Small *N*_e_*/N* demonstrates large variance in reproductive success (Akita, 2019; Wang, Santiago, & Caballero, 2016; Waples, 2016), resulting from the variance of reproductive potential (e.g., the big old fat fecund female fish hypothesis; Hixon, Johnson, & Sogard, 2014) or from the situation in which only some families successfully reproduce (referred to as the “Sweepstakes reproductive success” hypothesis, Hedgecock & Pudovkin, 2011). Moreover, if *N*_e_*/N* is invariant across years, then *N*_e_ may behave like an index of *N*, and vice versa (Luikart, Ryman, Tallmon, Schwartz, & Allendorf, 2010). However, if *N*_e_*/N* fluctuates across years, the trends can clarify the interpretation of environmental and/or anthropological factors, causing the variance of reproductive potential, family-correlated survivorship, or fluctuating population dynamics.

The estimation of *N*_e_*/N* has been performed by utilizing the estimated values of contemporary effective population size (*N*_e_) and census size (*N*), unless complete pedigree data and/or full census data are available. Additionally, there are numerous methods for estimating *N*_e_ from genetic markers (Wang et al., 2016, and the references contained therein). There are also numerous methods for estimating *N*, such as a mark-recapture method or population dynamics modeling with survey data (e.g., Kéry & Schaub, 2011; Methot & Wetzel, 2013; Quinn & Deriso, 1999; Seber, 1982). It is known that there are large variations in both estimators; thus, their combination (i.e., the estimator of *N*_e_*/N*) also shows large variation (Marandel et al., 2018; Palstra & Fraser, 2012). There is currently a little theoretical foundation for the estimator of *N*_e_*/N*, indicating no guidance for a sample size to ensure the required accuracy and precision.

Close-kin mark-recapture (CKMR) is a recently developed method for estimating *N* that utilizes the information about kinship in a sample. This was possible owing to the recent advances in genetic methods for kinship determination (Bravington, Grewe, & Davies, 2016; Bravington, Skaug, & Anderson, 2016; Hillary et al., 2018; Skaug, 2017) although similar methods have been proposed in the beginning of the 21st century (Nielsen, Mattila, Clapham, & Palsbøll, 2001; Pearse, Eckerman, Janzen, & Avise, 2001; Skaug, 2001). Besides, the rationale is that the presence of a kinship pair in the sample is analogous to the recapture of a marked individual in mark-recapture. Therefore, kinship pairs in the sample are less likely to be observed in larger populations; thus, the number of kinship pairs may reflect *N*. Since the original CKMR is designed to estimate adult abundance (i.e., *N*), the monitoring data for CKMR also produce the estimator of *N*_e_ by detecting half-sibling (HS) pairs within the same cohort (Akita, 2019). Since kinship determination is more accurate, this kinship-oriented estimation of *N*_e_ was presented in the context of the sibship assignment method (Wang, 2009) and is expected to provide a much more accurate estimator.

In this study, we propose a new method for estimating the ratio of contemporary effective mother size to the census size (*N*_e,m_*/N*_m_) in a population, as a proxy of *N*_e_*/N*. Assuming that kinships are genetically detected without any error, this method is based on the numbers of maternal-sibling (MS) and mother–offspring (MO) pairs in a sample. Thus, sampling is completed at a single breeding time; sampling offspring within the same cohort and mothers probably shares MO relationship with sampled offspring. Our model provides a nearly unbiased estimator of *N*_e,m_*/N*_m_ that explicitly incorporates two types of overdispersed reproduction (i.e., parental and nonparental variations; Akita, 2019), making it possible to target a species that shows iteroparity (i.e., multiple reproductive cycles during the lifetime) and/or sweepstakes reproductive success. First, we explain the modeling assumption and sampling scheme. Then, we analytically determine (nearly) the un-biased estimators of *N*_e,m_, 1*/N*_m_, and *N*_e,m_*/N*_m_, which are based on the numbers of MS and/or MO pairs. Finally, by running an individual-based model, we investigate the performance of the estimator and provide a guide for a sample size. Moreover, it is noteworthy that our modeling framework can be applied to diverse animal species. However, the description of the model focuses on fish species, which are presently the best candidate target of our proposed method.

## 2 THEORY

The main symbols used in this paper are summarized in Table 1.

**Table 1:**
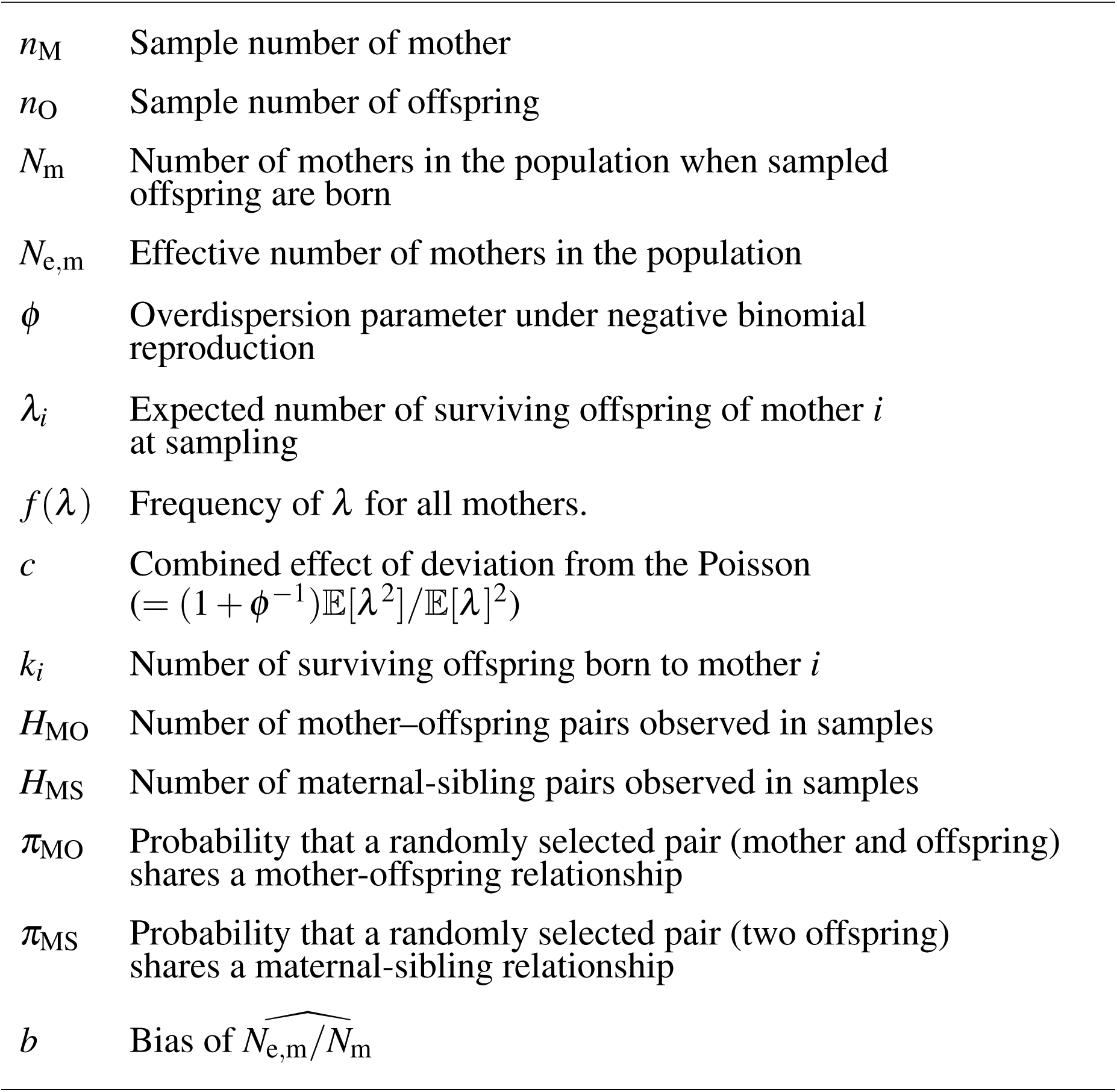
The list of mathematical symbols employed in the main text.

|  |  |
| --- | --- |
| $n_M$ | Sample number of mother |
| $n_O$ | Sample number of offspring |
| $N_m$ | Number of mothers in the population when sampled offspring are born |
| $N_{e,m}$ | Effective number of mothers in the population |
| $\phi$ | Overdispersion parameter under negative binomial reproduction |
| $\lambda_i$ | Expected number of surviving offspring of mother $i$ at sampling |
| $f(\lambda)$ | Frequency of $\lambda$ for all mothers. |
| $c$ | Combined effect of deviation from the Poisson<br>($= (1 + \phi^{-1})\mathbb{E}[\lambda^2]/\mathbb{E}[\lambda]^2$) |
| $k_i$ | Number of surviving offspring born to mother $i$ |
| $H_{MO}$ | Number of mother–offspring pairs observed in samples |
| $H_{MS}$ | Number of maternal-sibling pairs observed in samples |
| $\pi_{MO}$ | Probability that a randomly selected pair (mother and offspring) shares a mother-offspring relationship |
| $\pi_{MS}$ | Probability that a randomly selected pair (two offspring) shares a maternal-sibling relationship |
| $b$ | Bias of $\widehat{N_{e,m}}/N_m$ |

### 2.1 Hypothetical population

We suppose that there is a hypothetical population comprising *N*_m_ mothers and there is also no population subdivision or spatial structure. In this study, a mature female is called a mother even if she does not produce offspring. For mathematical tractability, we assume that only one spawning ground exists in which the mothers remain for the entire spawning season. Following Akita (2019), our modeling framework employs two types of overdispersed reproduction: parental and nonparental variations. Thus, the former indicates a variation caused by the mother’s covariates, such as age, weight, and residence time on the spawning ground, while the latter indicates a variation caused by non-random stochastic events during a series of reproductive episodes, which are independent of the mother’s covariates, such as family-correlated survivorship or the mating behavior effects (e.g., competition for males/females and correlation between mating opportunities of the mother and the number of her offspring). **Figure 1** illustrates a schematic diagram of the effects of parental and nonparental variations exemplified by age-dependent reproduction and family-correlated survival on kinship relationships in a population. Detailed definitions of parental and nonparental variations are stated in Akita (2019).

**FIGURE 1.**
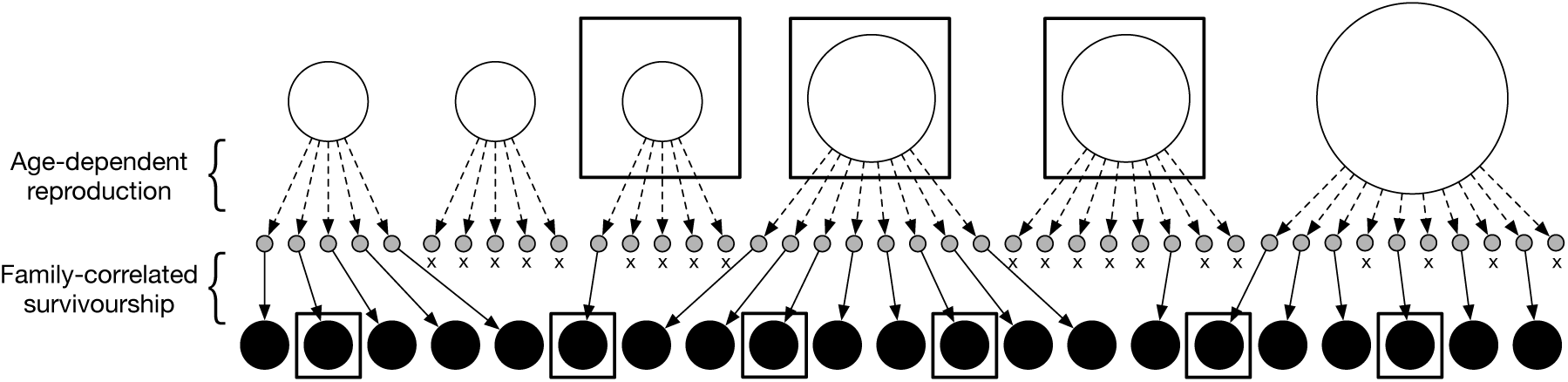
Example of relationships between mothers and their offspring number. The open, gray, and black circles represent mothers, their eggs, and their offspring, respectively. The area of an open circle indicates the degree of reproductive potential of each mother (i.e., *λ*_*i*_). The dotted and thin arrows denote mother–egg and egg–offspring relationships, respectively. The symbol x denotes a failure to survive at sampling. Sampled individuals are denoted with a bold line.

Let *k*_*i*_ denote the number of surviving offspring of mother *i* (*i* = 1, 2, *…, N*_m_) during sampling. It is noteworthy that *k*_*i*_ is assumed to be observed at the sampling, as explained in the next subsection. Following Akita (2019) and giving the expected number of the surviving offspring per mother during the sample timing, *λ*_*i*_ (> 0), *k*_*i*_ is assumed to follow a negative binomial distribution through a conventional parametrization:

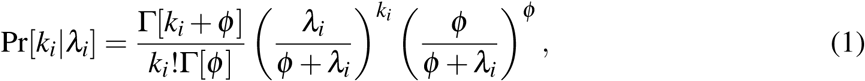

where *ϕ* (> 0) denotes the overdispersion parameter that describes the degree of the nonparental variation. Presently, *ϕ* is assumed to be constant across mothers, whereas the expected number of the surviving offspring (*λ*_*i*_) varies across mothers. The mean and variance of this distribution are denoted by *λ*_*i*_ and 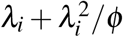, respectively. In the limit of infinite *ϕ*, this distribution becomes a Poisson distribution, which is given by 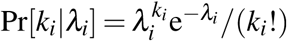. We assumed that *λ*_*i*_ is independent and identically distributed with the density function *f* (*λ*), which produces a parental variation. The shape of the density function is often complex, but it may be described by information, for example, the mother’s weight composition in the population. The specific form of *f* (*λ*) is given in **Supporting Information** and is used for running an individual-based model.

### 2.2 Sampling

To obtain the estimator of *N*_e,m_*/N*_m_, we utilize the number of MS and MO pairs observed in a sample. For the mathematical tractability, only one reproductive season is targeted for sampling. Thus, whether the sampling method does not affect our modeling framework whether it is invasive or noninvasive. In the population, *n*_M_ mothers are randomly sampled immediately after the end of the reproductive season. Additionally, in the population, *n*_O_ offspring are also randomly sampled but allowed to die before sampling. The numbers of MS and MO pairs are determined by pairwise comparison of all the sample individuals (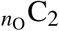 and *n*_M_*n*_O_ comparisons, respectively). As depicted in **Fig 1**, six offspring and three mothers are sampled and two MS and three MO pairs are observed.

In our modeling framework, if an MS pair also shares a paternal-sibling (PS) relationship, we count it as an MS pair and ignore the existing full-sibling (FS) relationship.

### 2.3 Effective mother size (*N*_e,m_) and the ratio to census size (*N*_e,m_*/N*_m_)

Akita (2019) derived the approximate probability showing that two offspring share an MS relationship with an arbitrary mother (denoted by *π*_MS_) as follows:

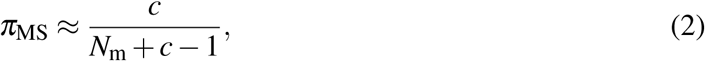

where

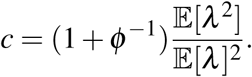

Without both parental and nonparental variations (i.e., *λ* is constant among mothers and *ϕ* → ∞), *π*_MS_ converges to 1*/N*, corresponding to the Poisson variance for all mothers in a population. Moreover, the effect of the two factors causing a deviation from the Poisson variance can be combined as parameter *c* (≥ 1). In the sequel, we refer to “overdispersion” as the distribution of the offspring number that results from this combined effect. By applying the combined effect, the variance of the offspring number can be given by

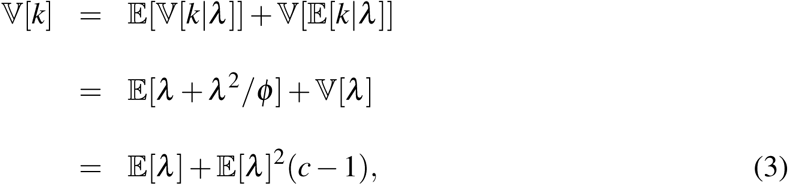

suggesting that the variance substantially increases with *c*.

Akita (2019) defined the contemporary effective mother size as follows:

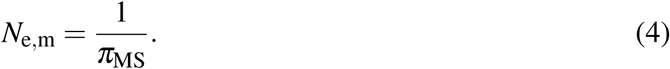

Besides, this definition is related to the inbreeding effective population size (Nordborg & Krone, 2002; Wang, 2009). When sampling from a single cohort in a population with overlapping generations, the effective mother size in our definition corresponds to the effective breeding mother size that produces a single cohort. We obtain the ratio of the effective mother size to census size using Eqs. 2 and 4 (Akita, 2019), and it is given by

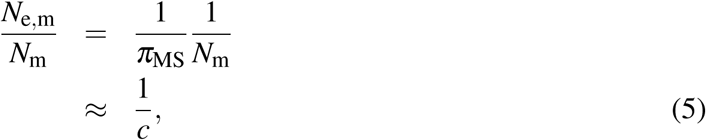

where *N*_m_ ≫ 1 is assumed for approximation.

### 2.4 Estimator of *N*_e,m_*/N*_m_

This subsection proposes the estimator of *N*_e,m_*/N*_m_ as follows:

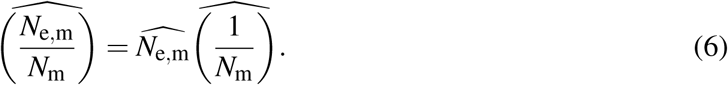

A “hat” denotes the estimator of a variable in this study. The requisite condition that satisfies Eq. 6 is independent of 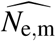 and 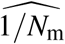 This property will be shown later in this subsection. Akita (2019) derived the nearly unbiased estimator of *N*_e,m_, which is given by

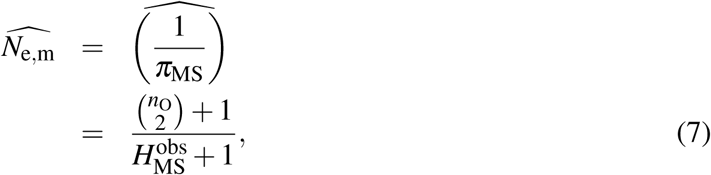

where 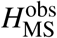 denotes the observed number of MS pairs in a sample. This estimator works well unless *n*_O_ is very small. Akita (2018) obtained a probability in which a randomly sampled mother and her offspring shares an MO relationship, which is given by

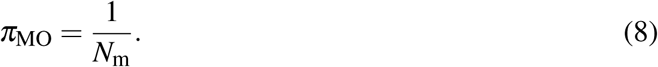

This indicates that *π*_MO_ is independent of the distribution of the offspring number. By definition of *π*_MO_, we can set its estimator by 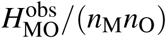, where 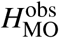 denotes the observed number of MO pairs in a sample. Thus, the estimator of 1*/N*_m_ can be determined as follows:

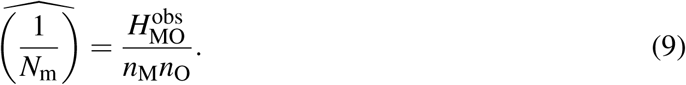

Finally, substituting 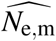 (Eq. 7) and 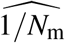 (Eq. 9) into Eq. 6, we obtain the estimator of *N*_e,m_*/N*_m_ as follows:

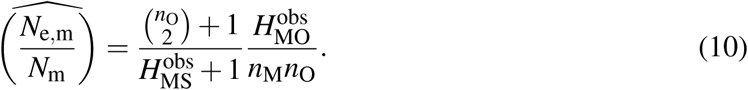

Let both *n*_M_ and *n*_O_ be given. We numerically confirmed that there is no correlation between 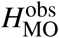 and 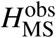 (results are not shown). To intuitively explain this independency, we consider three mothers (*i* = 1, 2, 3) and their offspring, and assume that (*k*_1_, *k*_2_, *k*_3_) = (3, 1, 1) and (*n*_M_, *n*_O_) = (1, 3). Moreover, when the three offspring born to the first mother are sampled (i.e., 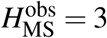), the expected number of MO relationship is one (= 1/3 × 3 + 1/3 × 0 + 1/3 × 0). Meanwhile, when an offspring is sampled from each mother’s offspring (i.e., 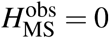), the expected number of MO relationship is also one (= 1/3 × 1 + 1/3 × 1 + 1/3 × 1). Therefore, we conclude that both 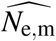 and 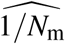 are independent, and 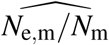 is expected to work well (see details in the **RESULTS** section).

The bias of 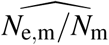 is defined by *b*, which is approximately given by (see **APPENDIX** for the derivation)

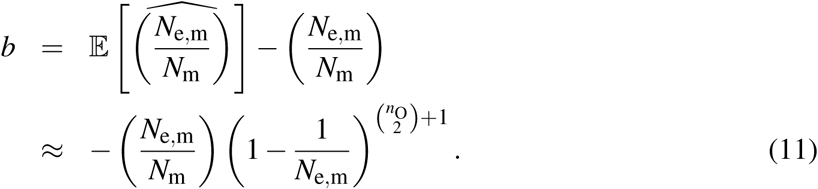

It is noteworthy that 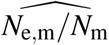 is downwardly biased, especially when *n*_O_ is very small. However, this bias may be ignored for a wide range of parameters (see details in the **RESULTS** section). Theoretically, *b* asymptotically converges to zero as *n*_O_ increases, making 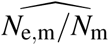 a nearly unbiased estimator. Moreover, as demonstrated in the **RESULTS** section, it is observed that an extremely skewed reproduction breaks down the unbiasedness (e.g., in the case that *c* = 20 and 100 in the results).

### 2.5 Individual-based model

We developed an individual-based model that tracks kinship relationships to evaluate the estimator’s performance (e.g., 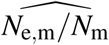). The population structure was assumed to be identical to that in the development of the estimators. In addition, the population comprised mothers and their offspring, and it was assumed to follow a Poisson or negative binomial reproduction. The expected number of the surviving offspring of a mother followed the density distribution *f* (*λ*) (see **Supporting Information** for details). We calculated overdispersion parameter (*c*) from *ϕ* and *f* (*λ*), as well as confirmed numerically that the value of *c* gives the same statistics of the estimators even if the combination of *ϕ* and *f* (*λ*) differs (results are not shown). Therefore, each offspring retained the mother’s ID, making it possible to trace an MS and MO relationship.

Let a parameter set (*n*_O_, *n*_M_, *N*_m_, *ϕ*, and parameters that determine *f* (*λ*)) be given. We simulated a population history in which *N*_m_ mothers generated offspring; this process was repeated 100 times. The sampling process for each history was repeated 10,000 times, acquiring 1,000,000 data points that were utilized to construct the distribution of the estimators for each parameter set. Furthermore, true value of *N*_e,m_ was calculated from *N*_m_ and *c* (Eqs. 2 and 4).

## 3 RESULT

We numerically evaluated the performance of 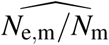 for the case in which the number of mothers, *N*_m_, and the combined effect of deviation from the Poisson, *c*, were unknown. Moreover, we changed the parameter values for *N*_m_ (10^3^ and 10^4^) and *c* (1, 10, 20, and 100). In addition, based on a given parameter set (*N*_m_ and *c*), we mainly addressed the number of samples (*n*_M_ and *n*_O_) required to obtain adequate accuracy and precision. In this study, we evaluated the performance of 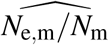 for specific ranges of the sample sizes (50-200 when *N*_m_ = 10^3^, and 200-1000 when *N*_m_ = 10^4^). Meanwhile, other estimators (i.e., 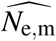 and 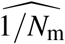) are also evaluated and provided in **Supporting Information**.

First, we evaluated the accuracy of estimators based on their relative bias calculated by applying the individual-based model, which is defined as follows: “(averaged estimator − true value)/true value.” For a given combination of *N*_m_ and *c*, the value of the relative error of 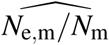 is represented on a heatmap as a function of *n*_M_ and *n*_O_, as dpicted in **Fig. 2**. For most of the investigated parameter sets, we observed that their relative error is less than 10%. As expected, the relative error is not affected by *n*_M_ since 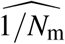 is exactly an unbiased estimator of 1*/N*_m_ (see Eq. A2 in **APPENDIX** and also **Fig. S2** in **Supporting Information**). Meanwhile, 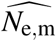 is downwardly biased when *n*_O_ is relatively small to true *N*_e,m_ (e.g., see *c* = 1 in **Fig. 2** and also **Fig. S1** in **Supporting Information**), as presented in Akita (2019); thus, 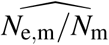 is downwardly biased. Contrary to the theoretical prediction for the direction of the bias (Eq. 11), relatively strong overdispersion results in an upwardly bias for 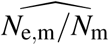 when *c* is relatively large (e.g., *c* = 20 and 100 in **Fig. 2a**). This inconsistency may be caused by the breakdown of the approximation for deriving 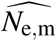 (Eq. S14 in Akita, 2019). Thus, as described in Eq. 3, extremely large *c* results in a large variance of offspring number, generating a situation in which the behavior of random variable *H*_MS_ far deviates from the binomial distribution.

**FIGURE 2.**
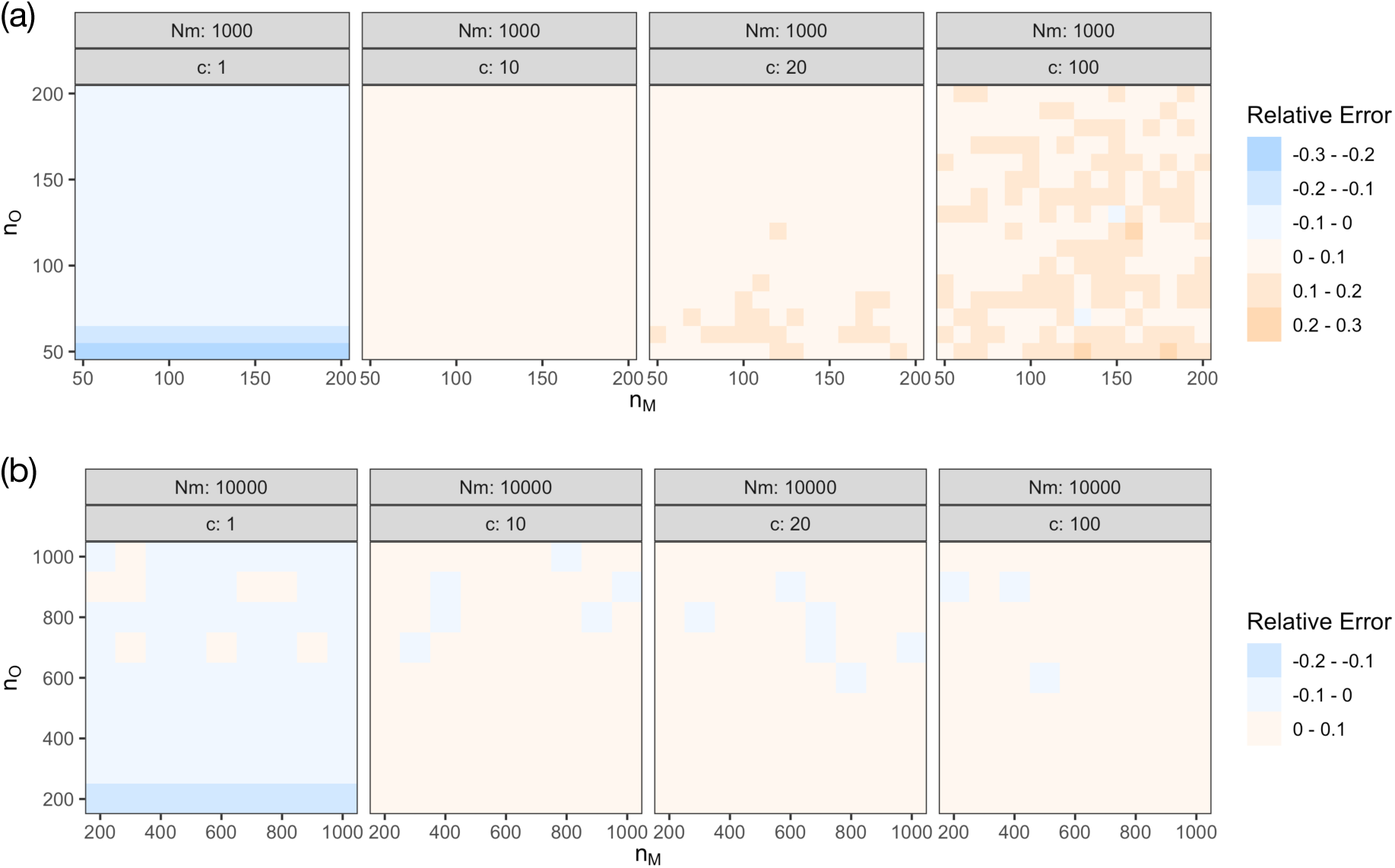
Heatmap showing the relative error of 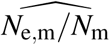 as a function of both *n*_M_ and *n*_O_: (a) *N*_m_ = 1,000, (b) *N*_m_ = 10,000.

Next, we evaluated the precision of estimators based on their coefficient of variation. As demonstrated in **Fig. 3**, the value of the coefficient of variation of 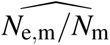 is also represented on a heatmap as a function of *n*_M_ and *n*_O_. For the investigated parameter sets, the degree of the coefficient of variation strongly depends on the sample sizes. As shown in **Figs. S3 and S4** in **Supporting Information**, the dependency results from the combined effects of variation of both 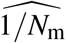 and 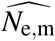. As *c* increases, it is noteworthy that the parameter space of sample sizes demonstrating large variation of 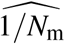 (e.g., *CV* > 30%) expands; however, when *c* is small (e.g., *c* = 1), relatively small *n*_O_ results in large variation of 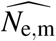 because of a relatively large *N*_e,m_.

**FIGURE 3.**
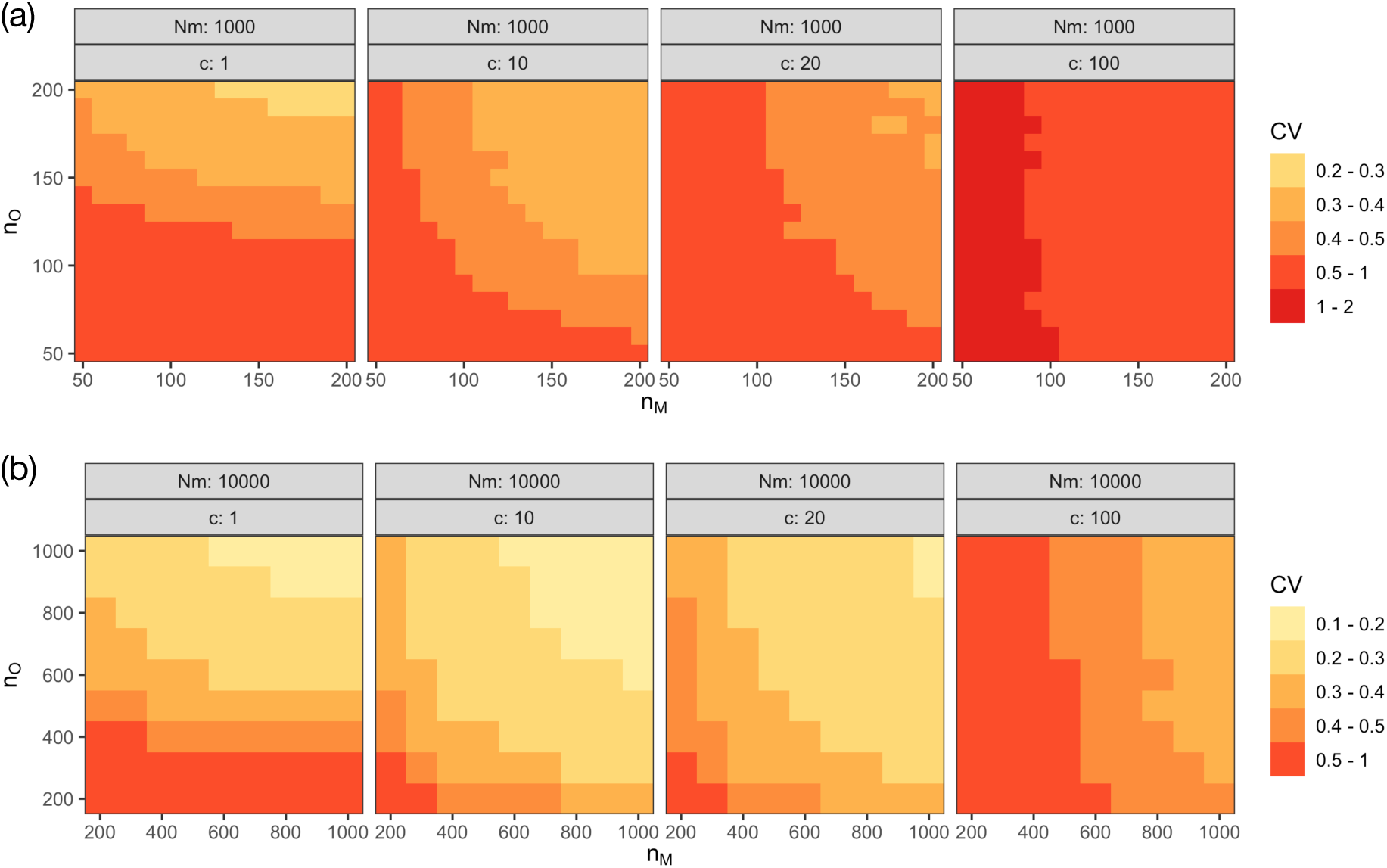
Heatmap showing the coefficient of variation of 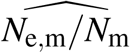 as a function of both *n*_M_ and *n*_O_: (a) *N*_m_ = 1,000, (b) *N*_m_ = 10,000.

## 4 DISCUSSION

We theoretically developed a nearly unbiased estimator of the ratio of contemporary effective mother size to the census size (*N*_e,m_*/N*_m_) in a population (Eq. 10). The proposed estimator is based on known MO relationship and MS relationships observed within the same cohort, in which sampled individuals in the cohort probably share MO relationships with sampled mothers (**Fig 1**). Moreover, the performance of the estimator (accuracy and precision) was quantitatively evaluated by running an individual-based model (**Figs. 2 and 3**). Meanwhile, the bias is analytically provided (Eq. 11). Our modeling framework utilizes two types of reproductive variations (Akita, 2019): variance of the average offspring number per mother (parental variation, denoted by *f* (*λ*)), and variance of the offspring number across mothers with the same reproductive potential (nonparental variation, denoted by *ϕ*). Additionally, these two effects result in a skewed distribution of offspring number and are summarized into one parameter (*c*) in the framework. Thus, our estimator can be calculated from sample sizes of mother and offspring (*n*_M_ and *n*_O_, respectively) and the observed numbers of MS and MO pairs (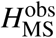 and 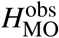, respectively), and it does not require other parameters. The rationale for this is the following: i) the frequency of MS and MO pairs contains information about *N*_e,m_ and *N*_m_, respectively; ii) the estimators of *N*_e,m_ and 1*/N*_m_ are independently determined based on a pedigree structure in the population and sample sizes, generating the estimator of *N*_e,m_*/N*_m_ by multiplying both estimators (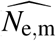 and 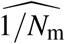). In this study, although 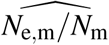 is considered as a proxy of 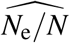, our theoretical results can easily be extended to the estimator of the ratio of contemporary effective father size to the census size if fathers are also sampled. The comparison of both ratios could clarify the underlying processes that differentiate between the sexes in the context of reproductive ecology.

The novelty of this study is that 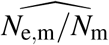 can be obtained only from the genetic data, and there are numerous advantages in using the proposed estimator instead of separately estimating *N*_e_ (via genetical method) and *N* (via non-genetical method). First, sampling and analyzing designs have become substantially simplified. Moreover, requirements for the proposed estimator are sampling of mothers and (potentially) their offspring in an appropriate time, and the extraction of their DNA that satisfies an adequate number of markers for kinship detection. In addition, both MO and MS pairs can be detected by a applying unified framework of genetic analyzes (there are many algorithms to detect kinship pairs from single nucleotide polymorphisms (SNPs) or short tandem repeats (STRs)), although an MS pair involves many more DNA markers (e.g., several thousands of SNPs are required for detection) than an MO pair (e.g., several hundreds of SNPs are required for detection). Second, our theoretical results guide sample sizes (*n*_M_ and *n*_O_) to ensure the required accuracy and precision, especially if the order of the number of effective mothers is approximately known. This is due to the simple formulation of the estimator determined only by the observed values (Eq. 10). Third, the proposed estimator directly reflects the amounts of *N*_e,m_ and *N*_m_ at the same timing (i.e., immediately after the end of the reproductive season), leading to a clear interpretation of the results, especially for genetic monitoring. For example, when the strong cohort is added to the spawning population in the beginning of the year, the estimator of *N*_e_ without reflecting this addition may results in an inappropriate estimation of *N*_e_*/N* (details of the temporal scale relevant to estimated *N*_e_ for each method were discussed in Wang et al., 2016).

Our modeling framework is presented by combining the context of the sibship assignment method (for estimating *N*_e,m_) and the CKMR method (for estimating 1*/N*_m_), which defines a kinship-oriented estimation of effective/census population size. Therefore, improvements to these methods directly contribute to the estimation of *N*_e_*/N*. Furthermore, the original theory of the sibship assignment method requires HS and FS pairs but does not require a distinction between the MS and PS pairs. This is a significant advantage due to the difficulty of the distinction from genetic data. However, the limitation of using MS or PS pair enables us to employ a nearly unbiased estimator of *N*_e_ for particular sex (Akita, 2019), which greatly improves the accuracy of the estimation of the *N*_e,m_ in this study and thus that of *N*_e,m_*/N*_m_. It is noteworthy that the estimator of 1*/N* is given by

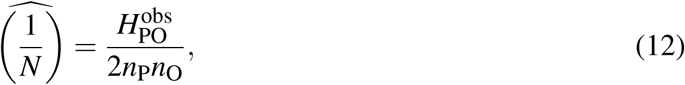

where *n*_P_ and 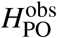 denotes the sample size of the parent and the observed number of parent–offspring (PO) pairs in a sample, respectively (Bravington, Skaug, & Anderson, 2016). The development of the unbiased estimator of *N*_e_ without a distinction between MS and PS pairs that could provide an unbiased estimator of *N*_e_*/N* coupled with Eq. 12, is a study for the future. Furthermore, using cross-cohort HS pairs, the CKMR method also provides the estimator of *N* (Bravington, Skaug, & Anderson, 2016) that does not require the sampling of the parent, which probably provides the estimator of *N*_e_*/N* only from unmatured samples. This perspective of the study will also be conducted in the future.

## Acknowledgments

The author thanks R. Nakamichi for fruitful discussions. This work was supported by JSPS KAKENHI Grant Number 19K06862.

## APPENDIX

### Derivation of the bias of 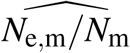

For calculation of the bias of 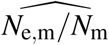, we require an expectation of the estimator given by

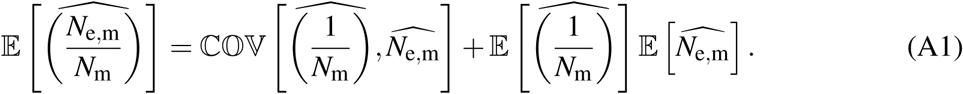

As stated in the main text, both 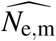 and 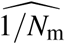 are independent. Thus, the first term in the right-hand side of Eq. A1 can be ignored. The expectation of 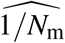 is given by

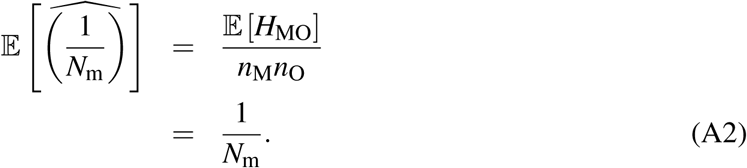

From the first to the second line of Eq. A2, we applied the relationship *π*_MO_ = 𝔼[*π*_MO_] = 𝔼[*H*_MO_]/(*n*_M_*n*_O_) and Eq. 8. Equation A2 indicates that 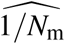 is the unbiased estimator. The expectation of 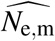 is given by

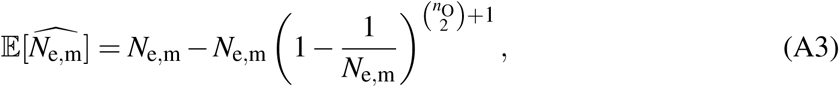

which is illustrated in Appendix D of Akita (2019). Together with these relationships, we can obtain the bias of 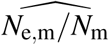 described in Eq. 11.

## Supporting Information

### Probability density function and its moment of *λ*

As stated in the main text, our modeling framework does not require the specific form of *f* (*λ*); it only requires the ratio of the second moment to the squared first moment (𝔼[*λ* ^2^]/𝔼[*λ*]^2^) instead. However, the specific form is required for evaluating the theoretical results (i.e., calculating the moment or running the individual-based model). Herein, we model an age-structured fish population that serves as a representative example, demonstrating both parental and nonparental variations. The following contents are almost the same as those of Akita (2019) except for the parameter values that produce the setting *c* = 20 and 100.

Suppose that the mean fecundity of a mother depends on her age. Let *λ*_*a*_ denote the mean fecundity, which is a function of age (denoted by *a*). The moment can be defined as 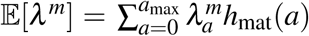, where *h*_mat_(*a*) is the frequency of mature mothers at a given age, and *a*_max_ denotes the maximum age. Thus, we can numerically obtain the moment by applying *λ*_*a*_ and *h*_mat_(*a*).

For marine species with a type-III survivorship curve, it is generally assumed that individual fecundity is proportional to weight. By utilizing the von Bertalanffy growth equation for body weight, *λ*_*a*_ is explicitly defined as a function of age as follows:

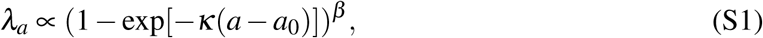

where *κ, a*_0_, and *β* are conventionally used parameters in the von Bertalanffy equation, and they denote the growth rate, the adjuster of the equation for the initial size of the animal, and the allometric growth parameter, respectively. To obtain a specific value of *λ*, a coefficient value of 10 multiplied by the right-hand side of Eq. S1 was used when running the individual-based model.

The frequency of mature mothers at a given age can be given as the following:

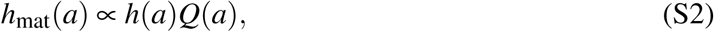

satisfying 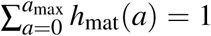, where *h*(*a*) and *Q*(*a*) denote the frequency and maturity at a given age, respectively. Although *f* (*a*) is affected by historical population dynamics and age-dependent survival, for simplicity, the mortality rate is assumed to be constant (i.e., age independent):

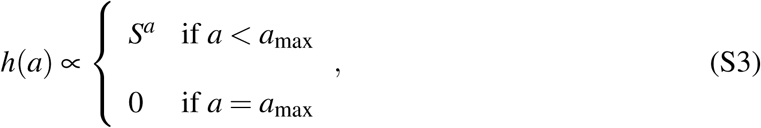

where *S* denotes a survival probability. The maturity at age (*Q*(*a*)) is assumed to be a knife-edge function, which is given by

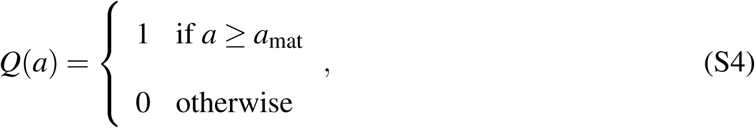

where *a*_mat_ denotes the mature age.

To calculate 𝔼[*λ* ^2^]/𝔼[*λ*]^2^, the required parameter set is (*a*_max_, *κ, a*_0_, *β, S, a*_mat_). In this study, for the purpose of representation, we fixed the values of several parameters as follows: *a*_max_ = 20, *κ* = 0.3, *a*_0_ = 0, *S* = 0.5 and *a*_mat_ = 0. In addition, we selected parameter value *c* (= (1 + *ϕ* ^−1^)𝔼[*λ* ^2^]/𝔼[*λ*]^2^) to be 1, 10, 20, and 100 for comparison with the results in the main text that are derived from the parameter set (*ϕ, β*) = (1000, 0.0009), (0.1302, 0.9), (0.06111, 0.9), and (0.01165, 0.9), respectively.

Finally, we provide specific forms of *f* (*λ*); thus, when *λ*_*a*_ and *h*_mat_(*a*) are obtained, *f* (*λ*) is given by

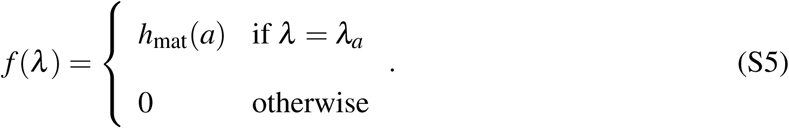

---

**FIGURE S1.**
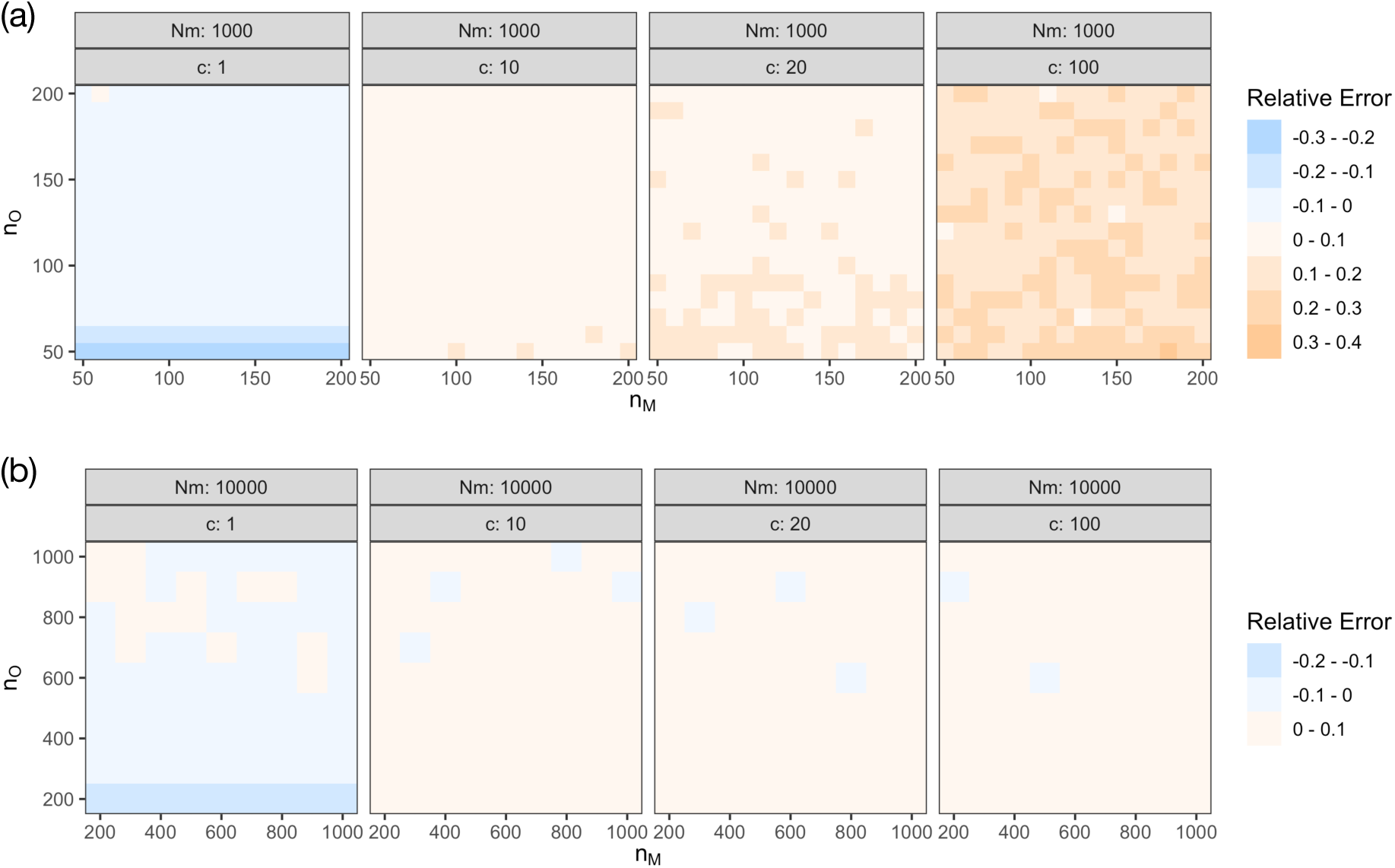
Heatmap showing the relative error of 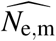 as a function of both *n*_M_ and *n*_O_: (a) *N*_m_ = 1,000, (b) *N*_m_ = 10,000.

**FIGURE S2.**
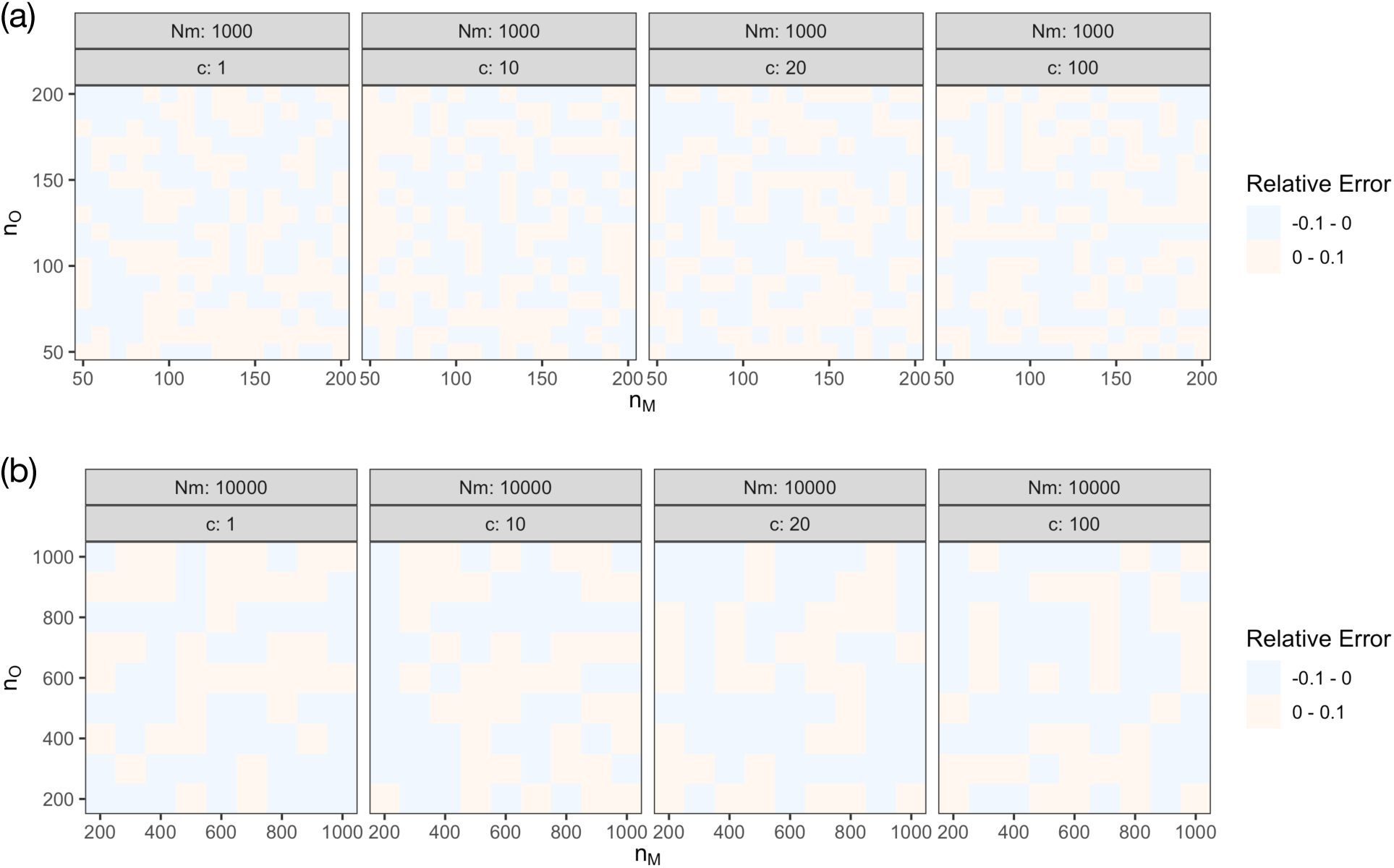
Heatmap showing the relative error of 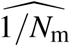 as a function of both *n*_M_ and *n*_O_: (a) *N*_m_ = 1,000, (b) *N*_m_ = 10,000.

**FIGURE S3.**
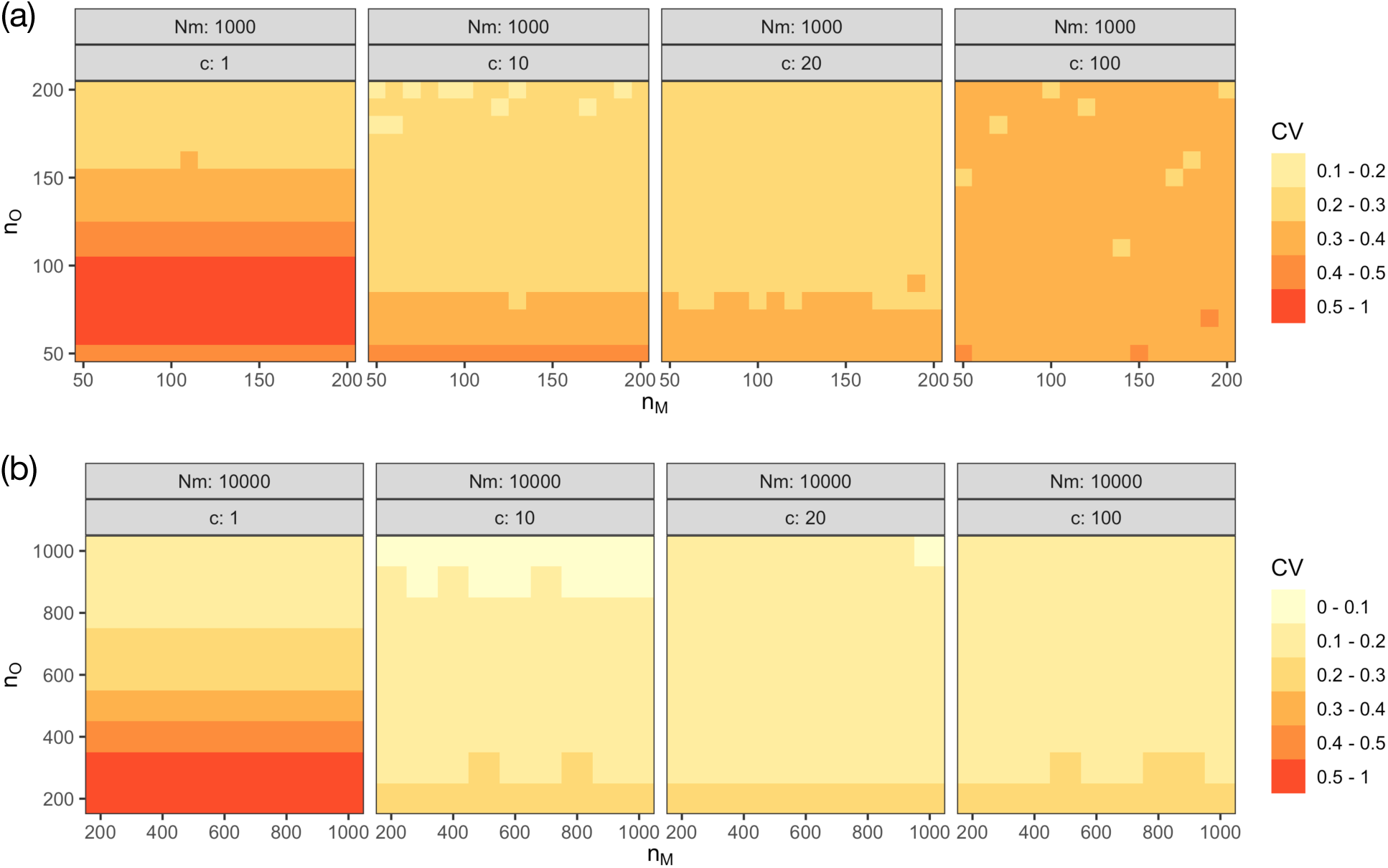
Heatmap showing the coefficient of variation of 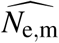 as a function of both *n*_M_ and *n*_O_: (a) *N*_m_ = 1,000, (b) *N*_m_ = 10,000.

**FIGURE S4.**
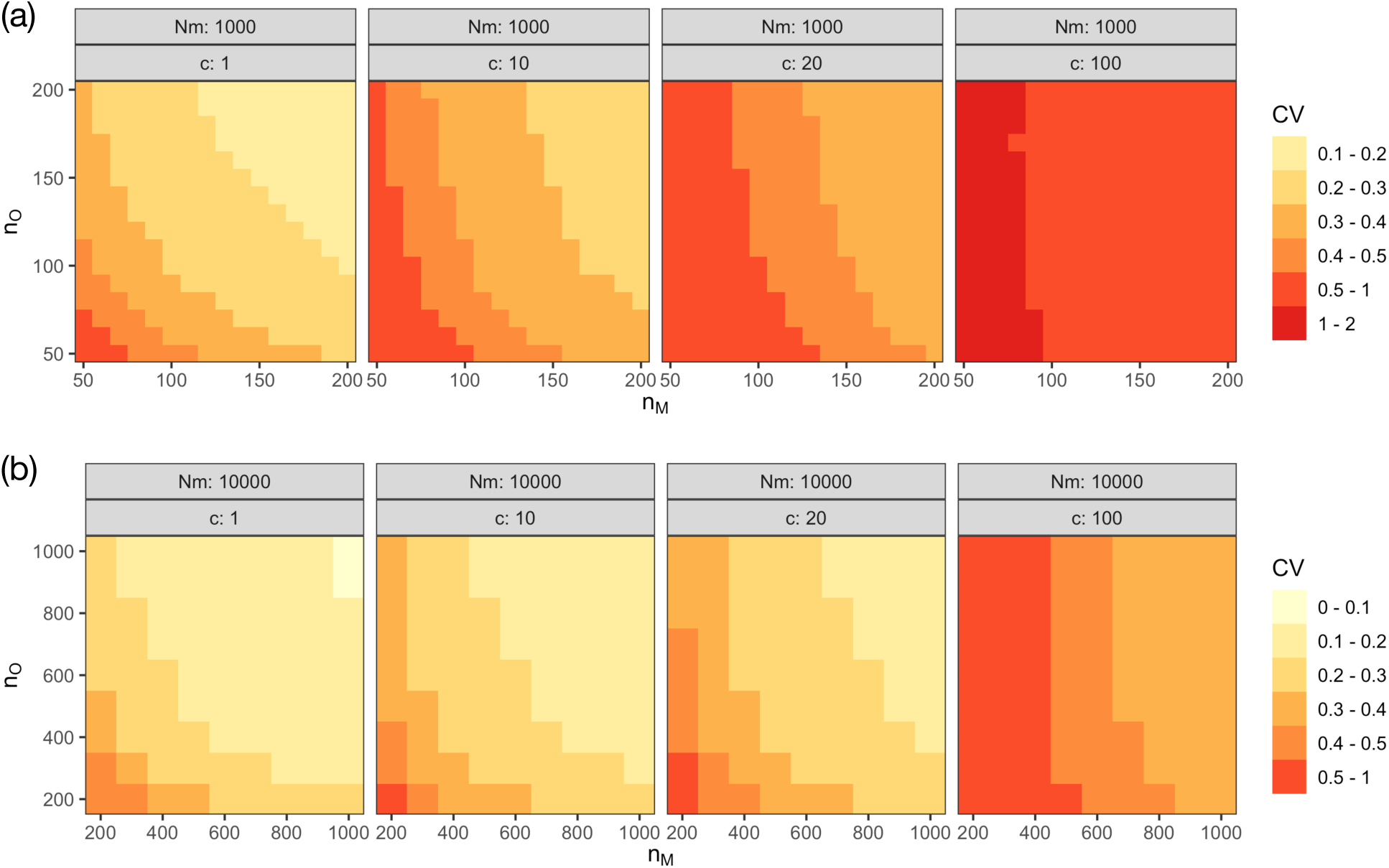
Heatmap showing the coefficient of variation of 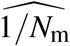 as a function of both *n*_M_ and *n*_O_: (a) *N*_m_ = 1,000, (b) *N*_m_ = 10,000.

